# Sound generation in zebrafish with Bio-Opto-Acoustics (BOA)

**DOI:** 10.1101/2020.06.09.143362

**Authors:** Itia A. Favre-Bulle, Michael A. Taylor, Emmanuel Marquez-Legorreta, Gilles Vanwalleghem, Rebecca E. Poulsen, Halina Rubinsztein-Dunlop, Ethan K. Scott

## Abstract

Hearing is a crucial sense in underwater environments for communication, hunting, attracting mates, and detecting predators. However, the tools currently used to study hearing are limited, as they cannot controllably stimulate specific parts of the auditory system. To date, the contributions of hearing organs have been identified through lesion experiments that inactivate an organ, but this makes it difficult to gauge the specific stimuli to which each organ is sensitive, or the ways in which inputs from multiple organs are combined during perception. Here, we introduce Bio-Opto-Acoustic (BOA) stimulation, using optical forces to generate localized sound in vivo, and demonstrate stimulation of the auditory system of zebrafish larvae with unprecedented control. We use a rapidly oscillated optical trap to generate vibrations in individual otolith organs that are perceived as sound, while adjacent otoliths are either left unstimulated or similarly stimulated with a second optical laser trap. The resulting brain-wide neural activity is characterized using fluorescent calcium indicators, thus linking each otolith organ to its individual neuronal network in a way that would be impossible using traditional sound delivery methods. The results reveal integration and cooperation of the utricular and saccular otoliths, which were previously described as having separate biological functions, during hearing.

---

Evolution has produced diverse approaches for hearing. Understanding different auditory systems in nature provides insights into the role of hearing in ecology, and has provided valuable design information for biomimetic microphone technologies[1, 2]. While mammalian hearing is based on a single organ, the cochlea, which only senses pressure waves, animals such as fish[3], crustaceans[4], and insects[5] all have multiple sensory organs that collectively provide hearing. In many cases, the precise functional contribution that each organ makes to auditory processing in these animals has remained elusive. This is because research in bioacoustics has traditionally been hampered by our inability to generate localized, intense sound in one organ without producing waves that propagate throughout all of the auditory sensory organs.

In fish, sound is sensed by otoliths, or “ear stones”, as well as the lateral line, which senses water flow across the body[3]. Since animals generally have a similar density to water, underwater sound travels almost unimpeded through their bodies, which makes it difficult or impossible to confine sound to specific sensory organs. Hearing research in aquatic animals therefore relies on speakers that stimulate all parts of the auditory system, with the relative contributions of different components only identified through destructive methods. For instance in fish, the specific contributions of the saccular and the utricular otoliths have only been tested by genetically or physically ablating the organs and comparing treated animals to controls [6-8]. The resulting models of fish hearing, which propose distinct and nonoverlapping roles for the different otoliths, have never been tested with direct and selective stimulation of the utricular and saccular systems.

Here, we present Bio-Opto-Acoustic (BOA) stimulation, in which optical forces generate sound to allow precisely controlled auditory stimulation of specific components of the auditory system. By generating the sound directly at each organ, we can localize the sound with a tight spatial resolution that is physically prohibited using propagating sound waves. Optical forces are applied using optical traps[9] (OT), which allow precise and non-invasive mechanical interactions. OT have been used for the manipulation of small transparent objects in a number of biological contexts[10-12], most notably in molecular biophysics[13, 14]. In this study, we have designed an optical system capable of applying BOA forces *in vivo* at frequencies ranging from 1Hz to 1kHz (Fig. 1a). We used this to vibrate the otoliths of larval zebrafish (Fig. 1b), allowing controlled stimulation of each auditory organ for the first time in any intact aquatic animal.

**Figure 1:**
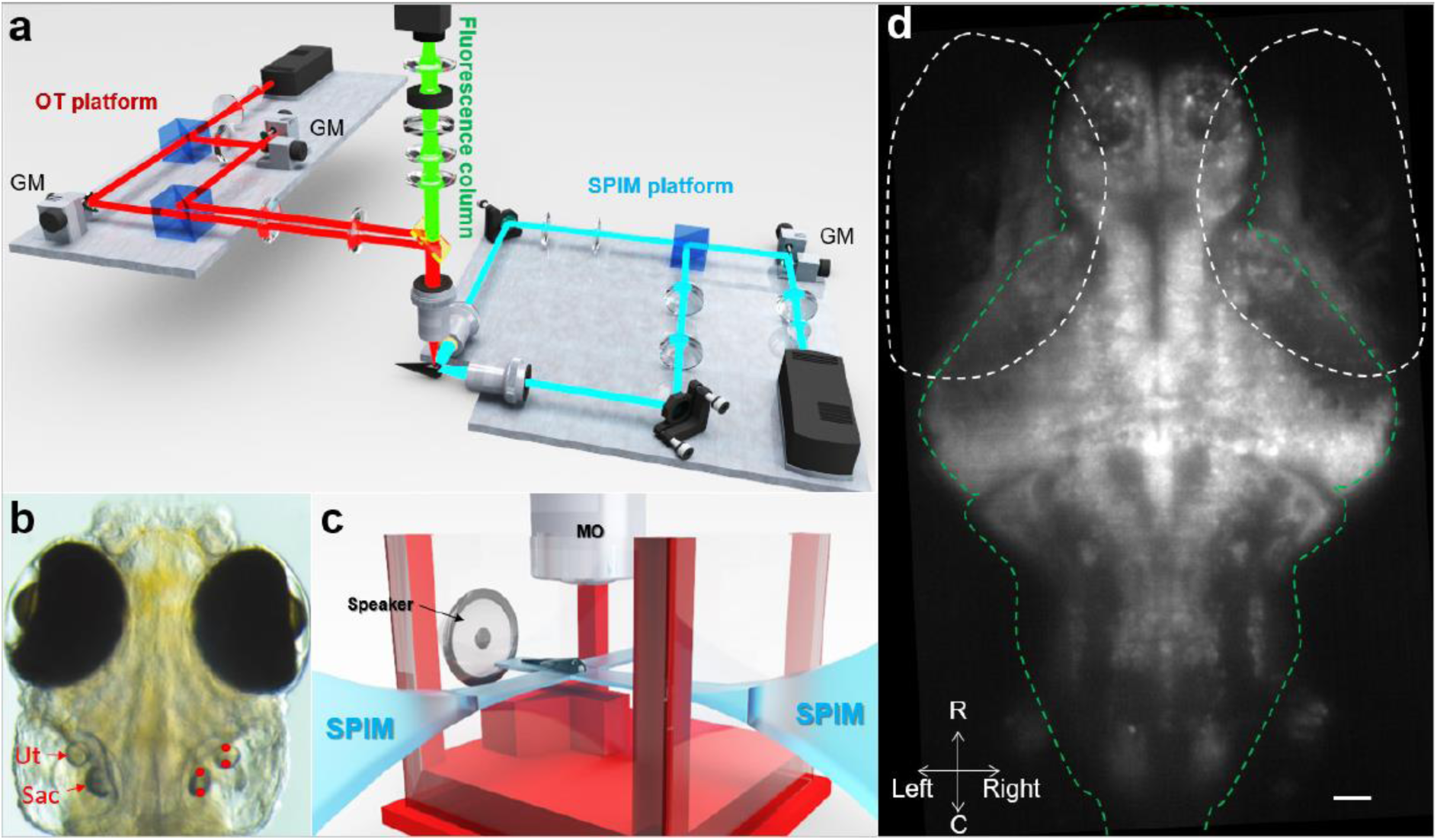
Optical and sound stimulation of the auditory system in zebrafish. **a**. Optical system comprising an OT platform for the generation of two optical traps, a SPIM platform for the illumination of a single plane of zebrafish brain, and a fluorescence column for the imaging of GCaMP6s emissions. Details can be found in the Methods. **b**. OT (red dots) were placed at two locations within the utricular (Ut) and saccular (Sac) otoliths. The galvo mirrors (GM) on the OT platform displaced each trap from one location to the other at a frequency ranging from 1Hz to 1 kHz. **c**. Sketch representing the placement of a larva in a custom built chamber, the SPIM planes, the microscope objective (MO), and the location of the speaker. **d**. Example of a fluorescence imaging frame recorded by the camera. The white dashed ovals indicate the eyes, and the green line delineates the brain. R, rostral; C, caudal. Scale bar indicates 10 μm.

The perception of BOA stimulation was verified by observing the brain’s responses to optical traps and to true sound delivered with a speaker. We combined both modes of stimulation with fluorescent calcium imaging of GCaMP6s in a light-sheet microscope, providing brain-wide recordings of the neuronal activity associated with the auditory and BOA stimuli (Fig. 1c,d). This allowed the targeted and systematic exploration of individual auditory organs and the contributions that they make to auditory and vestibular perception in this important neuroscience model system.

The bodies of fish, which have a density close to water, move in concert with traveling waves as sound passes through them. However, their otoliths, mostly made out of a tightly packed calcium carbonate crystal, are much denser than water. Consequently, otoliths within fish do not move as much as the body during auditory stimulation. This relative motion between the otoliths and the rest of the fish causes deflection of hair cells in the ear, thus producing a neural signal that feeds into auditory processing circuitry[15]. Acceleration sensed at low frequency is referred to as vestibular, while higher frequencies that are in our own hearing range are referred to as auditory, with a distinction between the two modalities typically drawn at approximately 10Hz. Past studies have suggested that the two types of otolith in larval zebrafish, the utricle and saccule, sense vestibular and auditory stimulation respectively[16-20], but the historical inability to stimulate these organs independently limits the strength of this interpretation.

Using the optical system presented in Figure 1a, we applied alternating OT to opposite sides of the otoliths at various frequencies, creating oscillations of the targeted otolith. These vibrations of the otoliths relative to the fish body are similar to what a sound wave would produce (Fig. 2a,b).

**Figure 2:**
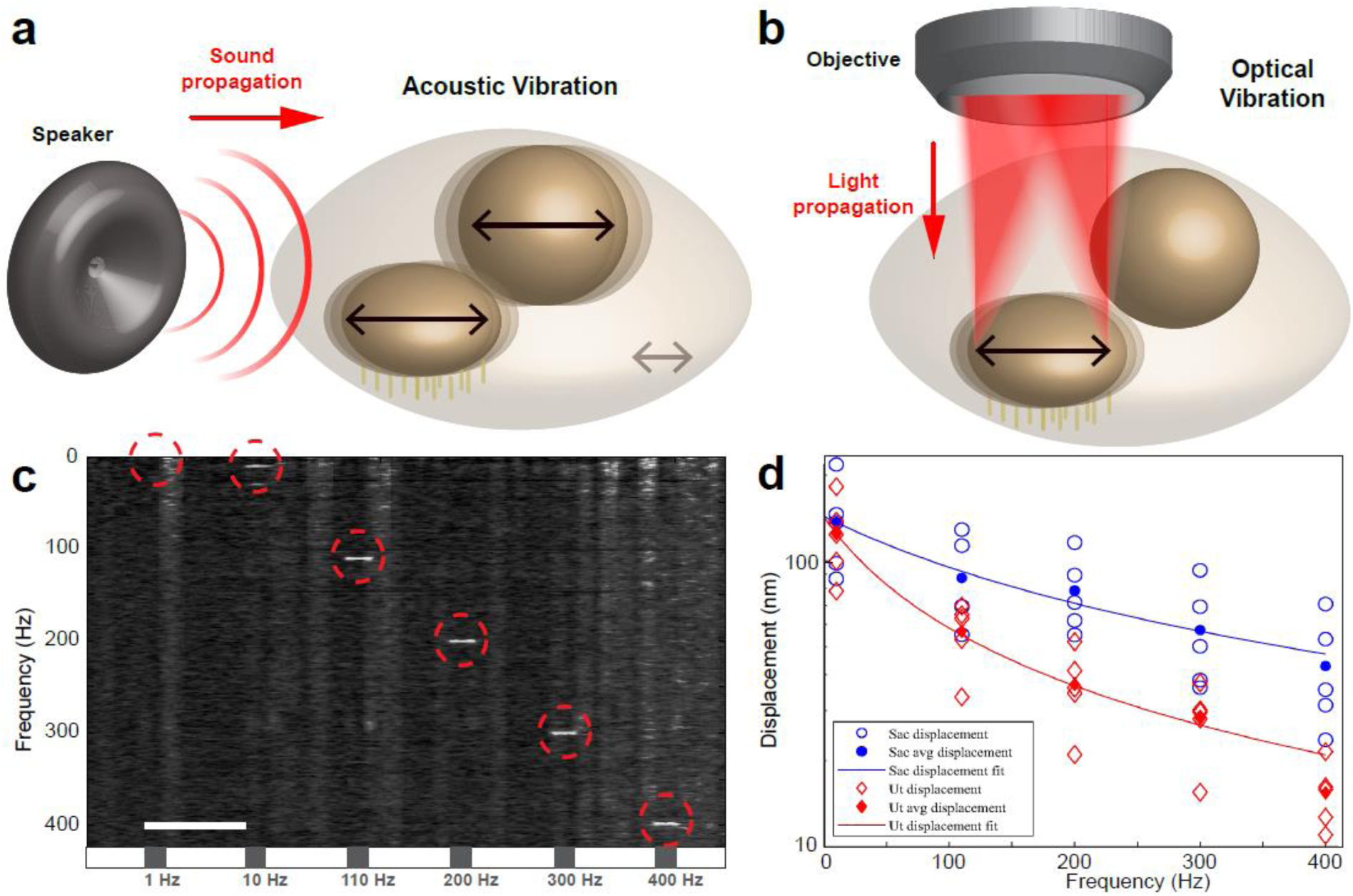
Mechanics of sound perception in zebrafish. **a**. Schematic illustration of sound propagation from a speaker and the resulting movements of the otoliths in larval zebrafish ear. **b**. Illustration of OT targeting from one side of one otolith to the other at high speed, and the resulting selective vibration of the targeted otolith. **c**. Average spectrogram (normalized over frequency) of the position of the optically manipulated saccule and utricle over time across 5 fish. Grey boxes on the timeline represent 1s of OT stimulus. The number written under the box represents the OT frequency of the stimulation. See Sup Fig. 1 for more details on the movements of each type of otolith. Scale bar 5s. **d**. Measurements of otolith displacements at different frequencies of BOA stimulation, displayed on a logarithmic scale. Fit was performed to Eq. (1), with fitting parameters describing *γ* and *k* and neglecting mass.

The motion of each otolith under either auditory or BOA stimulation can be expressed through Newton’s law: *F* = *ma*. In the case of auditory stimulation, it is the acceleration of the otolith relative to the body that is relevant, which results in an effective acceleration of *a*_*sound*_ = (1 − *ρ*_*f*_/*ρ*_*ot*_)*a*. We see that the high mass density of the otolith *ρ*_*ot*_ compared to the ear fluid *ρ*_*f*_ is essential for high sensitivity. Optical forces can replicate acoustic stimulation by generating similar accelerations of the otolith.

When solving Newton’s law equation in the Fourier domain, the solution of the otolith position can be expressed as (details in Supplementary Information):

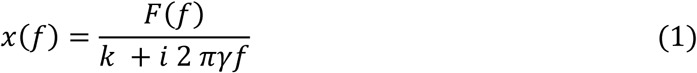

Where *γ* is the viscous drag coefficient of ear fluid, k the elasticity constant of hair cells, and where the total force *F* can include optical force, acoustic sounds, and background forces such as the heartbeat, blood flow, muscle movement, and thermal fluctuations. At low frequencies we simply expect x∼*F*/*k* limited by the hair cell stiffness. At higher frequency we anticipate the oscillation amplitude to scale as 1/*f*.

To quantify the amplitudes of these vibrations at different frequencies, we measured the otoliths’ displacement during BOA stimulation (See Methods and Supplementary Figure 1). Vibrations of each otolith can be visualized and quantified from the spectrogram of its displacement over time (Fig. 2c). While displacement at low frequencies (1 Hz) are masked by other movements in the living animal (heartbeat, blood flow, and others), displacements at higher frequencies can be identified and quantified (red circles in Figure 2c).

The results confirm the displacement of otoliths using BOA stimulation in vivo. At 10 Hz, we measured displacements of roughly 140 nm, while at 400 Hz, those displacements decreased to about 50 nm for the saccular otolith and 15 nm for the utricular otolith (Fig. 2d) with 400 mW laser power. As expected from Eq. (1) the amplitude of motion decreased with higher frequency. This is unsurprising, since temporally shorter OT forces should produce smaller displacements of the otolith. We found larger displacements across the frequency range for the saccular otolith (Fig. 2d), in spite of its larger size and mass, likely owing to a more favourable geometry that produces stronger trapping forces during BOA stimulation.

The range of frequencies in our BOA stimulus train (Supplementary Figure 2.a) include those associated with vestibular stimuli (1Hz), those at the interface of vestibular and auditory stimuli (10Hz), and those that are considered to be in the auditory range (100 and 1000Hz). As such, we can flexibly test specific otoliths for their ability to detect and relay information to the vestibular and auditory systems in a way that is not possible with real-world auditory and vestibular stimuli, observing the resulting sensory responses by combining our BOA stimulation with whole-brain calcium imaging in stationary larvae.

Our modified microscope (Fig. 1) was used to perform brain-wide volumetric GCaMP imaging (using the *elav3:H2B-GCaMP6s* transgenic line[21], expressing GCaMP6s in the nuclei of all neurons) at 4Hz volumetric imaging rate (details in Methods). This approach was used to map brain-wide responses to a range of BOA frequencies, along with 100 Hz auditory tones. An example fluorescence image is shown in Figure 1d. We first automatically segmented regions of interest (ROIs) generally corresponding to individual neurons across the brain, and then extracted signals through time for each ROI, using CaImAn package[22], as stimuli were presented (See Methods). We then performed a linear regression to identify ROIs responsive to the auditory tones and BOA stimuli, followed by a k-means clustering to identify classes of ROIs (clusters) with distinct response profiles to the stimuli. Following this step, we selected clusters responsive to auditory tones and that were consistently represented across all six larvae tested (see Supplementary Figure 2 and the selection criteria detailed in the Methods). Finally, we warped the 3D structures of all six animals’ brains onto one another and onto the Z-brain atlas of the larval zebrafish brain[23], providing a registered reference brain for our responses, and mapped each responsive ROI back to its 3D position within the brain (See Methods). This approach allowed us to identify and locate all ROIs responsive to tones or BOA stimulation, and to register them within a common reference.

Our goal was to identify clusters responsive to tones and to compare our targeted BOA stimulation of the utricle, the saccule, or both to actual auditory stimulation that would affect both otoliths. To remove the complication arising from unilateral versus bilateral stimulation, we pierced the left ear of each larva, rendering it deaf to auditory stimulation. This ensured comparable stimulation of the right ear only as we applied both BOA and auditory stimuli at 100 Hz.

We observed that the clusters responsive to 100 Hz tones from the speaker were also responsive to 100Hz OT stimulation, confirming that BOA stimulation taps into natural auditory circuits in the brain (Fig. 3.a). Interestingly, cluster 1 shows that the simultaneous trapping of the utricle and saccule enhance the neuronal response in a super-additive manner (Fig. 3b). This suggests that the utricle contributes to the detection of a wide range of frequencies. We additionally found a saccule-specific category of ROIs (cluster 2). These ROIs show no pronounced preference for higher frequencies, responding across the range of 1Hz to 100Hz (Fig. 3c), which surprisingly suggests the involvement of the saccular otolith in the detection of low frequencies that are more vestibular than auditory. These data indicate that both the utricle and the saccule contribute to detecting a wide range of frequencies rather than processing distinct frequency ranges in parallel, and further suggest that auditory and vestibular perception in larval zebrafish are mediated by both otoliths in concert rather than each being carried out by a designated otolith.

**Figure 3:**
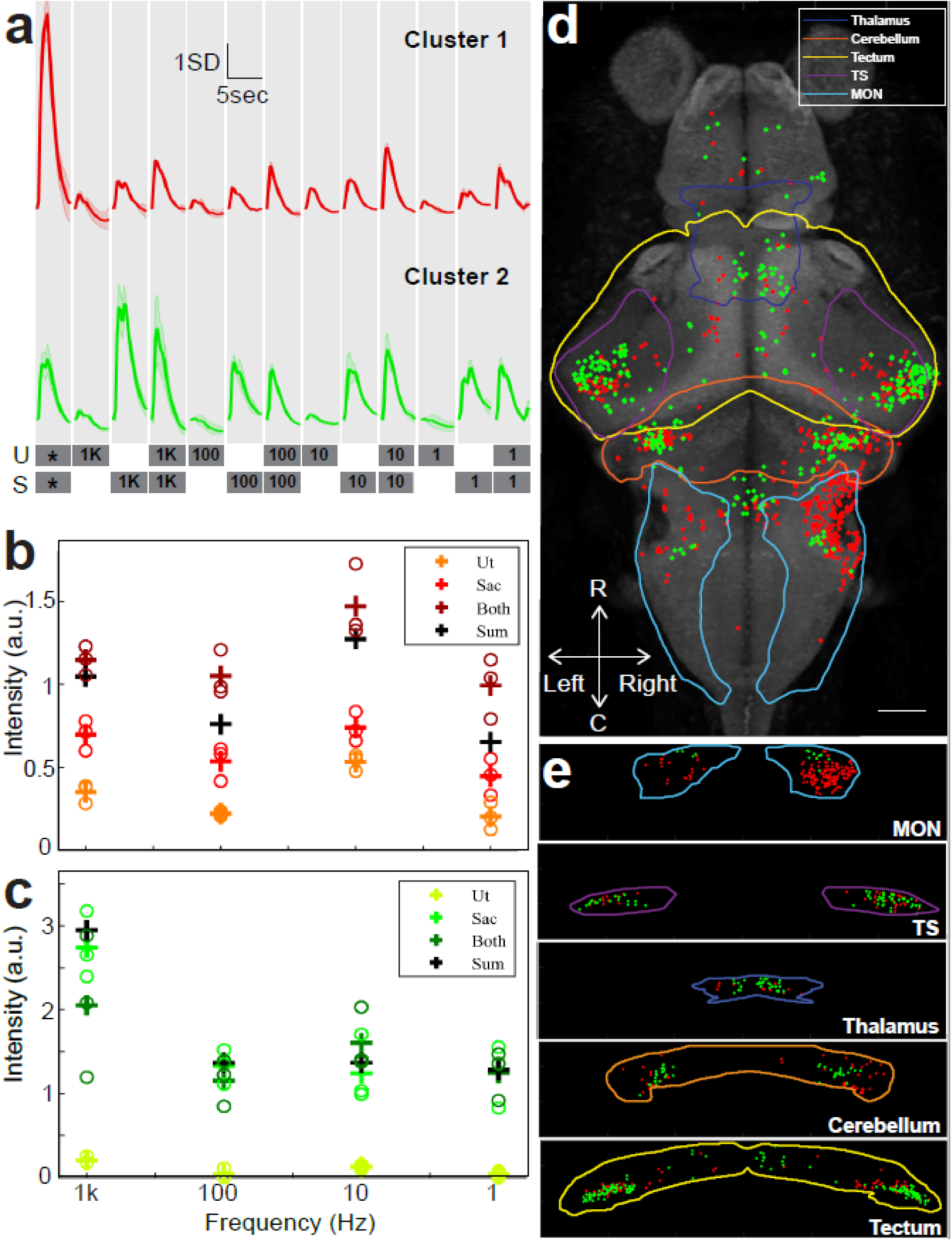
Responses distribution. **a**. Average profile of 100Hz responsive clusters during auditory and BOA stimulation. The two bottom lines detail the stimulus train. Grey boxes specify the stimulus windows (1sec of stimulation and 4sec of rest) and the otolith targeted (U, utricle; S, saccule). Numbers on the grey boxes specify the frequency of BOA stimulation in Hz. * represents a 100Hz auditory tone from a speaker. **b, c**. Intensity of responses of cluster 1 **(b)** to and cluster 2 **(c)** to BOA on the utricle, saccule, and both at variable frequencies. The sum of utricle and saccule responses is indicated in black. **d**. Locations of ROIs belonging to two functional clusters of auditory neurons, viewed dorsally. Data are from six fish. Scale bars 50μm. **e**. Distributions of each cluster in the medial octavolateralis nucleus (MON), torus semicircularis (TS), thalamus, cerebellum and tectum, rotated to produce a coronal view.

In terms of their distributions across the brain (Fig. 3d), these responses occur in structures of the primary auditory pathway, including the medial octavolateralis nucleus (MON) in the hindbrain, the torus semicircularis (TS) in the midbrain, and the dorsal thalamus in the forebrain (Fig. 3e). These responses are similar to what has been previously found in auditory studies with zebrafish larvae, which, along with matching to the auditory stimuli in our own study, corroborates the efficacy of the BOA as a proxy for auditory stimulation[24, 25].

The MON is the first relay center in the brain for auditory and vestibular information[26], and given that the left ear is inactivated and BOA is applied to the right ear, responses to both auditory and BOA stimulation are preferentially found in the right MON (Fig. 3d, e). Activity in the contralateral (left) MON is presumed to result from connections between the left and right MON, as have been observed for auditory responses in other fish species[27, 28] and for lateral line information in zebrafish[29]. We note that this pronounced asymmetry is lost in subsequent processing centers, suggesting that later stages of auditory and vestibular processing are performed bilaterally in this system.

Apart from the structures of the main ascending auditory pathway, the BOA stimulation also elicits responses in the cerebellum and tectum (Fig. 3e). The activity of the cerebellum could be modulating the auditory responses[30], or could be related to motor responses[25]. On the other hand, the tectum is known to integrate multiple sensory modalities, including auditory information[31]. It is possible that auditory information in the tectum can be used to generate a spatial map, as the homologous superior colliculus does in mammals[32].

Finally, we note a transition from the MON where we find a preponderance of cluster 1 ROIs, to brain regions later in the auditory pathway where we see more balance between the clusters or a greater number of cluster 2 ROIs. This suggests a possible shift from general responses in the MON (with responses to all tested frequencies and drawn from both otoliths) to more selective responses (favoring high frequencies, and with selective control by the saccular otolith) later in the auditory processing pathway.

In this study, we have introduced BOA as a method for the precise stimulation of the auditory system with light. Our observations, while they provide the first unambiguous accounting of saccular and utricular contributions to brain-wide auditory processing, likely understate the breadth and complexity of these sensory pathways. Further analyses with bilateral BOA stimulation, a greater number of stimulus frequencies, and a correspondingly greater diversity in functional clusters, will be necessary fully to map the nuances of each otolith’s contribution to auditory and vestibular processing. Such studies will allow the detailed characterization of frequency discrimination, analyses of the convergence and differentiation of vestibular and auditory pathways in the brain, and the ontogeny of auditory and vestibular processing during development. As these structures are part of the main ascending auditory pathway of teleost[30, 33], which are shared with amphibians and mammals[30, 34-36], BOA represents an important new tool for the exploration of the vertebrate auditory system.

## Methods

### Animals

All procedures were performed with approval from The University of Queensland Animal Welfare Unit (in accordance with approval SBMS/378/16). Zebrafish (*Danio rerio*) larvae, of either sex, were maintained at 28.5°C on a 14 hr ON/10 hr OFF light cycle. Adult fish were maintained, fed, and mated as previously described[37]. All experiments were carried out in nacre mutant *elavl3:H2B-GCaMP6s* larvae[21] of the TL strain.

### Sample preparation

Zebrafish larvae at 5 days post-fertilization (dpf) were immobilized in 2% low melting point agarose (LMA) (Sigma-Aldrich) on microscope slides. Using the thin sharp end of a pulled pipette, the left ears of zebrafish larvae were pierced. The fish were released from LMA and placed back in their incubator. The following day, these larvae were immobilized dorsal side up in 2% LMA on microscope slides. Each embedded fish was transferred to custom made, 3D printed chamber[38], which was filled with E3 media[37]. Larvae were then allowed to acclimate for 15 min prior to imaging on the custom-built dual optical trapping microscope presented in Figure 1.

### Dual Optical Trapping system and targeting

The dual Optical Trapping (OT) system (Figure 1) was composed of an infrared laser (1070nm IPG Photonics YLD-5 fiber laser), a half wave plate that rotates polarization by 45 degrees, and a polarizing beam splitter that splits the incoming beam into two beams of the same intensity with perpendicular polarizations. The two independent beams (Trap 1 for the utricle and Trap 2 for saccule) were reflected off a galvo mirror (Thorlabs GVSM002/M). The two beams were recombined with a second polarizing beam splitter and a telescope (150 mm and 300 mm focal length). An additional a lens (150 mm focal length) was added into Trap 2 path to displace this trap +20 µm in Z (above Trap 1) in order to reach the Saccule, located around 20 µm above the utricle. The beams were then reflected off a 950 nm cut-off wavelength shortpass dichroic mirror in the imaging column, and projected onto the back focal plane of a 20x 1NA Olympus microscope objective (XLUMPLFLN-W). This created two tightly focused spots at the imaging plane of the microscope objective. The positions (x,y) of each pair of spots (two spots for each otolith) were steered with the galvo mirrors. The two galvo mirrors were driven with Arduinos (Leonardo) in order to place the OT beam at precise (x,y) locations, and oscillated between these predetermined locations at variable frequencies (1, 10, 100, and 1000 Hz). Two shutters (Thorlabs SHB1T) allowed independent gating of the OT and were also driven using an additional Arduino (Leonardo). A laser power of 400 mW was used and gauged using a power meter at the focal plane of the 20x 1NA objective. The optimal targeting position within each utricular and saccular otolith was determined by the position of the beam in the otoliths that produced maximum of applied force, which was found to result from traps positioned between 1 and 3 µm from the edge of the otolith[39]. Once the trapping positions were optimised, we performed the calcium imaging or the otolith vibration imaging as described below.

### Imaging and analysis of otolith movements

In order to image both the utricular and saccular otoliths at high speed (1kHz), we built a system to provide trans illumination of the fish with a bright white LED light under the specimen. Using μManager[40], we cropped the video recordings to a tight region around each otolith to allow the acquisition frame rate to reach 1kHz. Each otolith was recorded separately, and otolith motion was estimated from recorded movies using an efficient subpixel image registration by cross-correlation method[41]. Using this method, we calculated the displacement of each otolith in X and Y in response to the optical manipulation. Since the resulting traces were noisy, we represented the data in the Fourier domain using a spectrogram (Figure 2 and Supplementary Figure 1).

### Fluorescence imaging system

Calcium imaging and OT were performed through the same 20x objective. For the fluorescence imaging of a chosen depth on the PCO edge 5.5 camera, a combination of filter (Thorlabs FF01-517/520-25), tube lens (180 mm focal length, Thorlabs AC508-180-A), relay lenses (Thorlabs AC254-125-A-ML), ETL (Optotune EL-10-30-Ci-VIS-LD driven with Gardasoft TR-CL180) and offset lens (Eksma Optics 112-0127E) was constructed as described by Fahrbach et al.[42]. The scanning light sheet was generated using a 488nm laser (OBIS 488 lx), scanned with 2D galvo mirrors (Thorlabs GVSM002/M), a 50/50 beamsplitter, and a 1D line diffuser (RPC Photonics EDL-20-07337), as previously described[43].

With this configuration, we were able to scan 250 µm of brain tissue above the original imaging plane, where utricle otolith is placed. One galvo scanning direction (Y) created the light sheet while the second direction (Z) created the depth scan in the sample. The two mirrors were driven independently using Arduinos (DUE) with custom-written code. The Y scanning was a sawtooth scan at 600 Hz, which was synchronized to the camera acquisition to ensure similar illumination for each camera acquisition. The Z galvo was driven in 10 ms steps to scan the light-sheet in Z through the sample. The 50/50 beamsplitter created two light sheets, one projecting into the rostral side and one into the right side of the fish. The 1D line diffuser was placed just after the galvos to reduce shadowing effects in the planes[43]. The imaging system was controlled using μManager, based on ImageJ[44, 45]. In our experiments, an exposure time of 10 ms was chosen for each plane during volumetric imaging, with laser power output of 60 mW, which was attenuated to 1.5 mW for each plane at the sample. A total depth of 250 µm in Z was scanned[46], with 25 planes at 10 µm intervals, resulting in a 4 Hz volumetric acquisition rate. We commenced laser scanning 30s prior to imaging neural activity to eliminate responses to the onset of this off-target visual stimulus.

### Extraction of fluorescent traces

The volumetric scan was first transformed into a hyperstack in Fiji[47], and then separated into individual slices. We used the CaImAn package to analyze our images[48, 49] and extract the fluorescent traces of each ROI from every slice (http://github.com/flatironinstitute/CaImAn). We compensated for mall movements using a rigid registration [50]. The greedy roi method was used to initialize, for each slice, 4000 components from which to extract, demix, and denoise the fluorescent traces using an autoregressive model of order 1. [48]. We used a correlation threshold of 0.8 to merge overlapping ROIs and avoid over-segmentation. The components were updated before and after the merge steps, empty components were discarded, and the components were ranked for fitness as in[48].

### Whole brain analysis of fluorescent traces

The active ROIs and their respective fluorescent traces were further analyzed in MATLAB with a custom-written code, as previously described[51]. The traces from six fish were pooled and z-scored. Regressors were built for the stimulus train presented with a typical GCaMP response at each stimulus onset (Supplementary Figure 1a.). A linear regression was performed between all the fluorescent traces and the regressors. The coefficient of determination (r^2^) of the linear regression models was used to select stimulus responsive ROIs, and we chose a 0.2 threshold based on the r^2^ distribution of our models to allow for conservative filtering of the data (Supplementary Figure 1b.). The next step was clustering with k-means method. The fluorescent traces passing the linear regression test were clustered into 120 clusters using k-means with the cityblock distance and five replicates. All the clusters’ averages were correlated with the regressors, and the clusters responsive to tones and specific to the optical vibration of the saccule with a response at least 1SD above baseline were selected. The fluorescence traces within each resulting cluster were compared to the regressors using linear regression and the ROIs showing r2 values above 0.4 were selected (Supplementary Figure 2c.).

Finally, clusters were filtered with the following selection criteria:

1. Responsivity to each tone stimuli as a GCaMP6s profile,
2. Responsivity to each tone stimuli with a response above 1SD to base line,
3. Less than 90% of the ROIs within the cluster are represented in a single fish.

### Spatial registration of fluorescence imaging to a reference brain

We used Advanced Normalization Tools (ANTs, https://github.com/ANTsX/ANTs) to compute the diffeomorphic map between the time-averaged 3D image stack of each fish and the H2B-RFP reference of Z-brain[23, 52, 53]. The same mapping was used to warp the centroid coordinates for each ROI of interest to the H2B-RFP frame of reference, which includes 294 segmented brain regions [23]. We used MATLAB to represent each ROI centroid as a sphere within the Zbrain reference brain image.

## Acknowledgments

Support was provided by an NHMRC Project Grant (APP1066887), a Simons Foundation Pilot Award (399432), a Simons Foundation Research Award (625793), and two ARC Discovery Project Grants (DP140102036 & DP110103612) to E.K.S. Support was provided by ARC Discovery Project Grants (DP180101002) to H.R. Support was also provided by the Australian National Fabrication Facility (ANFF), QLD node.

## Author contributions

I.A.F. and E.K.S. designed the project. I.A.F. and M.A.T. built the optical system. I.A.F. and

R.E.P. designed the fish chamber. I.A.F. carried out the experiments. I.A.F., E.M-L and G.V. processed the fluorescent neuronal data. I.A.F., M.A.T, E.M-L, G.V., H.R. and E.K.S wrote the manuscript.

