## Supplementary material for "Sound generation in zebrafish with Bio-Opto-Acoustics (BOA)"

### Supplementary Information

Itia A. Favre-Bulle, Michael A. Taylor, Emmanuel Marquez-Legorreta, Gilles Vanwalleghem, Rebecca E. Poulsen, Halina Rubinsztein-Dunlop, Ethan K. Scott

#### Modelling of Otolith motion under periodic stimulation

The motion of each otolith undergoing sound pressure or BOA stimulation can be studied analytically using Newton's law:

$$\sum \vec{F} = m\vec{a}$$

The forces present in the direction of the BOA stimulation when the otolith is manipulated are the viscous drag from the ear fluid $F_D$, the pulling forces of the hair cells $F_{HC}$ on the otolith and a force $F$ which can include optical force, acoustic sounds, and background forces such as heartbeat, blood flow, muscle movement, and thermal fluctuations.

$$F(t) + F_D(t) + F_{HC}(t) = ma$$

Under the assumption that the hair cells are elastic, and that we apply optical forces and sound pressure in the elastic regime of hair cells, we can deduce from Hooke's law that $F_{HC}$ can be expressed as $F_{HC} = -kx$, where $k$ is the spring constant and $x$ is the otolith displacement. Therefore:

$$m\frac{d^2x}{dt^2} + \gamma\frac{dx}{dt} + k\,x(t) = F(t)$$

Where $\gamma$ is the viscous drag coefficient, and m is the mass of the otolith. This second order differential equation can be solved in the Fourier domain, which would give us[1]:

$$x(f) = \frac{F(f)}{k\; + i\,2\,\pi\gamma f - m(2\,\pi\,f)^2}$$

At low frequencies we have x~F/k. The displacement amplitude is limited by the hair cell stiffness. At higher frequency we anticipate viscous drag to dominate, which causes the oscillation amplitude to scale as 1/f. Inertial motion becomes significant at high frequency (f>γ/m), but the damping is expected to be sufficiently large that inertial motion is not significant at auditory frequencies. Therefore, we can approximate the otolith displacement x in the Fourier domain to be:

$$x(f) = \frac{F(f)}{k\; + i\,2\,\pi\gamma f}$$

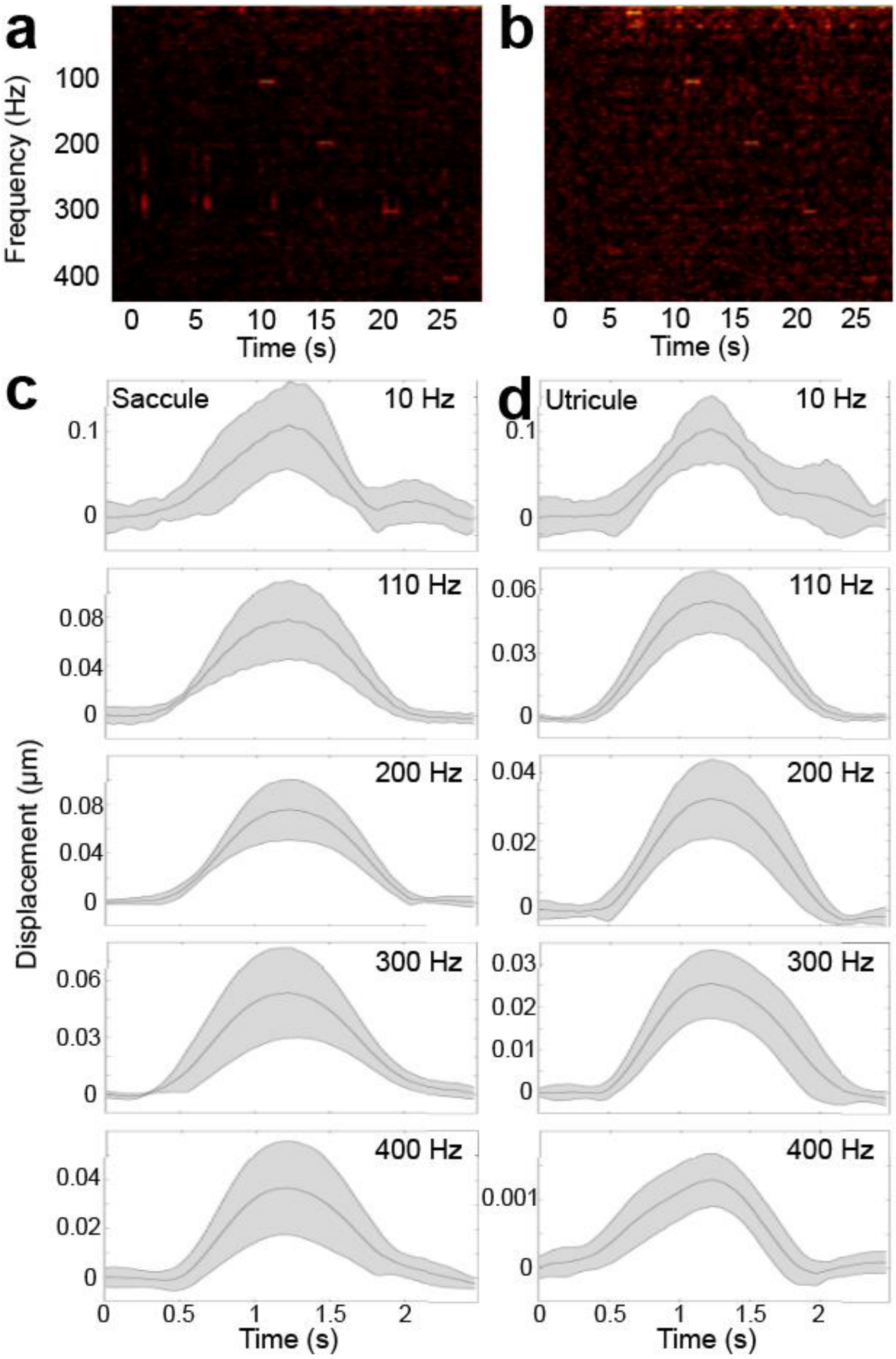


**Supplemental Figure 1. Detecting otolith movements at specific frequencies under optical trapping. a.** Example spectrogram representing motion of the saccular otolith in one fish with 1s window. **b.** A similar spectrogram showing utricular motion in one fish with 1s window. **c, d**. Average curves (solid line) and standard deviation (grey area) across 5 fish of the amplitude envelopes at specific frequencies (10, 110, 200, 300 and 400) detected for movements of the saccular **(c)**, and utricular **(d)** otoliths under optical manipulation. Each curve corresponds to one line of the spectrogram, truncated around the time of the stimulus.

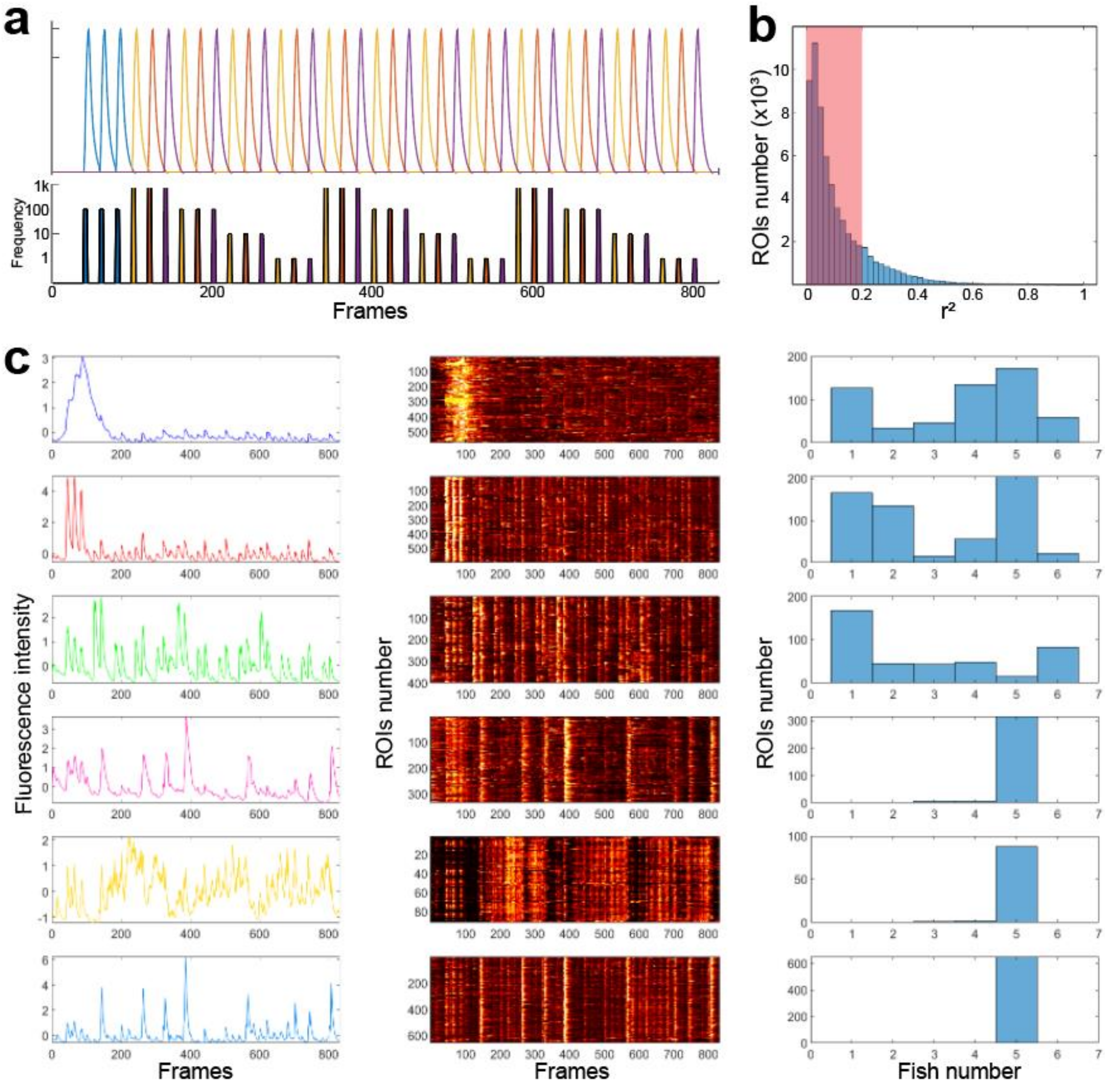


**Supplemental Figure 2. Clusters filtering. a.** Top line: regressors used for the linear regression step. Blue corresponds to the auditory tone, yellow to trapping of the utricular otolith, red to trapping of the saccular otolith, and purple to simultaneous trapping of both otoliths. Bottom line: stimulus train at various frequencies, with colors coded as above. **b.** Histogram of $r^2$ values showing the 0.2 threshold applied to all ROIs after the linear regression step. **c**. All clusters with average responses to tones > 1SD. First column shows the average traces of all ROIs in the cluster, and the second column shows raster plots of the individual ROIs' responses. Third column shows the number of ROIs drawn from each of the seven fish in the dataset. The first, fourth, fifth, and sixth clusters failed to meet the exclusion criteria (see Methods), and were not analyzed further. Cluster 1 did not pass selection criterion 1, while clusters 4, 5 and 6 did not pass selection criterion 3.
